# One-for-all gene inactivation via PAM-independent base editing in bacteria

**DOI:** 10.1101/2024.06.17.599441

**Authors:** Xin Li, Ying Wei, Shu-Yan Wang, Shu-Guang Wang, Peng-Fei Xia

## Abstract

Base editing is preferable for bacterial gene inactivation without generating double strand breaks, requiring homology recombination or highly efficient DNA delivery capability. However, the potential of base editing is limited by the adjoined dependence on the editing window and protospacer adjacent motif (PAM). Herein, we report an unconstrained base editing system to enable the inactivation of any genes of interests (GOIs) in bacteria. We first employed a dCas9 derivative, dSpRY, as the effector to build a base editor with activation-induced cytidine deaminase, releasing the dependence on PAM. Then, we programmed the base editor to exclude the START codon of a GOI instead of introducing STOP codons to obtain a universal approach for gene inactivation, namely XSTART, with an overall efficiency approaching 100%. By using XSTART, we successfully manipulated the amino acid metabolisms in *Escherichia coli*, generating glutamine, arginine, and aspartate auxotrophic strains. The effectiveness of XSTART was also demonstrated in probiotic *E. coli* Nissle 1917 and photoautotrophic cyanobacterium *Synechococcus elongatus*, illustrating its potential in reprogramming clinically and industrially relevant chassis. To be noticed, we observed a relatively high frequency of off-target events as a trade-off for the efficacy and universality.

## Introduction

Gene inactivation is essential for biological research and innovations. In bacteria, a predominant strategy is to disrupt the coding sequences (CDS), which usually deploys a “dead-or-alive” selection. For the conventional knock-in method, bacteria have to integrate an antibiotic resistance gene into a certain genomic locus to survive the corresponding antibiotics, thus leading to the disruption of a CDS (Xia et al., 2019). This has been excelled by the revolutionary CRISPR-Cas system. The RNA-guided Cas nuclease finds a specific genomic locus and generates a double-strand break (DSB). When a repairing template (donor DNA) is supplied, the DSB will be repaired through homologous recombination (HR), resulting in living cells with designed and clean edits in the genome (Knott and Doudna, 2018). Otherwise, the bacteria will die. Despite these advantages, the toxicity of Cas proteins, the low HR activity, and the requirement for efficient DNA delivery are making CRISPR-Cas-based gene inactivation challenging in bacteria (Collias et al., 2023; Vento et al., 2019; Yu et al., 2022).

Base editing deploys the CRISPR-Cas system and deamination for cytosine-to-thymine (C-to-T) or adenine-to-guanine (A-to-G) transitions in the genome at a single-nucleotide resolution (Gaudelli et al., 2017; Komor et al., 2016; Nishida et al., 2016). By using base editing, premature STOP codons can be introduced to genes of interest (GOIs) for gene inactivation (Kuscu et al., 2017). Notably, base editing employs a nuclease-deactivated Cas (dCas) protein with lower toxicity to bacteria, and it does not demand on HR for the desired editing, eventually relieving the reliance on highly efficient DNA delivery strategies to enable abundant transformants surviving from a “dead-or-alive” selection (Collias et al., 2023; Xia et al., 2020). These advantages empower base editing a preferable tool for gene editing, and its capacities have been demonstrated in various bacteria, including *Escherichia coli* (Banno et al., 2018), cyanobacteria (Lee et al., 2023; Wang et al., 2023), acetogens (Xia et al., 2020) and marine bacteria (Wei et al., 2023).

Two intrinsic features, however, constrain the potential of base editing. First, as a CRISPR-Cas-based system, it relies on a specific protospacer adjacent motif (PAM). Second, it has a specific hot-spot editing window, within which highly efficient base editing can be achieved. For instance, a base editor employing dCas9 from *Streptococcus pyogenes* and the activation-induced cytidine deaminase (AID) from *Petromyzon marinus* recognizes an NGG PAM and the hot-spot editing window lies between positions −16 to − 19 in a spacer (the nucleotide next to the PAM as position −1) (Rees and Liu, 2018; Wang et al., 2021). In addition, a premature STOP codon can only be introduced based on four codons (Wang et al., 2021). Taking together, to inactivate a GOI, we must find a specific codon (e.g., CAG coding for Gln) within a certain editing window (positions −16 to −19) of a spacer adjacent to a particular PAM (e.g., NGG), where the spacer is preferably located in the first half of the CDS to avoid unexpected truncations. Relieving these constraints would substantially enhance and expand the utility of base editing.

Here, we designed and established a one-for-all gene inactivation strategy for bacteria with unconstrained base editing. First, we employed a dCas9 derivative, dSpRY (Christie et al., 2023; Walton et al., 2020), as the effector to build a cytosine base editor with AID, releasing the dependence on PAM. Then, we programmed the base editor to exclude the START codon (e.g., ATG and GTG) of a GOI instead of introducing STOP codons, namely XSTART. By using XSTART, we successfully manipulated the amino acid metabolisms in *E. coli*, generating glutamine, arginine, and aspartate auxotrophic strains. The capability of XSTART was also demonstrated in probiotic *E. coli* Nissle 1917 (EcN) and photoautotrophic cyanobacteria.

## Results

### Design of a PAM-independent base editor for bacteria

SpRY is an engineered Cas9 derivative, finding a protospacer without the requirement of PAM, while a preference for NRN (R stands for A or G) over NYN (Y stands for T or C) PAM was reported (**Figure 1A**) (Walton et al., 2020). We first modularly assembled the dSpRY (D10A and H840A) and AID from *P. marinus* as the editing module with uracil DNA glycosylase inhibitor (UGI) and Leu-Val-Ala (LVA) degradation tag to enhance the performance. Then, the guide RNA (gRNA) was implemented. Finally, we chose a temperature-sensitive replication origin for selecting, maintaining, and rescuing the working plasmids **(Figure S1A)**. Presumably, the resulting dSpRY-AID system would release its requirement of PAM, and with such a system, we can relocate the target loci to the hot-spot editing window of the base editor for improved editing efficiency.

**Figure 1.**
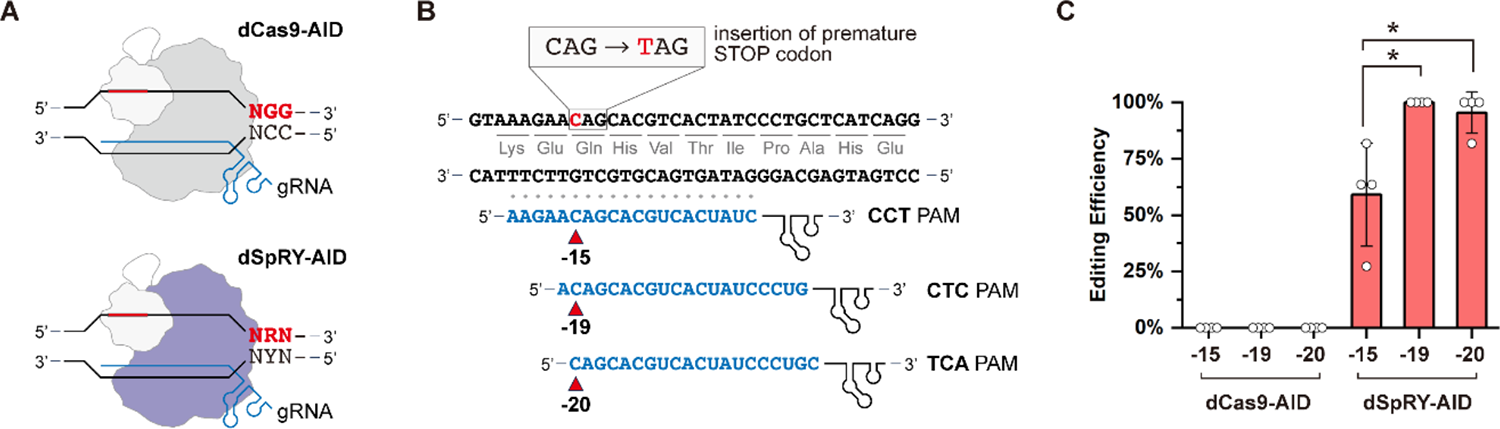
Unconstrained base editing with dSpRY. **(A)** Schematic illustration of dCas9-AID and dSpRY-AID mediated base editing, where dCas9-AID recognizes a NGG PAM and dSpRY-AID recognizes a NRN or NYN PAM. R stands for A or G, and Y stands for T or C. **(B)** Design of the three spacers targeting *glnA* based on dSpRY-AID editing module, where the targeted C was relocated in different editing windows with unconstrained PAM. The red arrows indicate the designed editing loci. **(C)** Base editing efficiencies of dCas9-AID and dSpRY-AID systems. The results represent the means of four biologically independent replicates and the error bars indicate the standard deviation (SD). The differences were statistically evaluated by *t*-test (\**p* < 0.05).

To demonstrate the system, we chose *glnA*, encoding the glutamine synthetase, as a target, and attempted to insert a premature STOP codon in Q30 (Figure 1B). We designed three gRNAs **(Table S3)** with the target locus located in positions −15, −19, and −20, respectively, with corresponding PAMs CCT, CTC, and TCA (Figure 1B). Meanwhile, a dCas9-AID system was generated for comparison with the same gRNAs **(Figure S1B)**. As expected, the dCas9-AID system showed no editing efficacy with these non-NGG PAMs (Figure 1C). To the contrary, the dSpRY-AID system showed promising activity, even when the target nucleotide was located in position −15. We observed that when the loci of the target moved to positions −19 and −20, the editing efficiencies significantly increased reaching 100% and 95.45 ± 9.09%, indicating the capacity of dSpRY-AID for improving editing performance (Figure 1C). To be noticed, we also found mixed signals in the resulting colonies at the first round of selection, which is a common issue for base editing (Wei et al., 2023; Xia et al., 2020), while strains with pure edits could be obtained via one more round of segregation **(Figure S2)**.

### One-for-all gene inactivation by excluding the START codon (XSTART)

Only four specific codons can be converted to the three STOP codons, but the START codons, which intrinsically contain the nucleotide G, are universal and can be eliminated with merely one C-to-T transition on the non-coding strand (Figure 2A). The resulting DNA sequence has, if any, limited influences as translation will not initiate anymore. Therefore, we designed XSTART for gene inactivation by excluding the START codons with dSpRY-AID. Theoretically, XSTART can edit any GOI by designing a gRNA targeting the non-coding strand with CAT or CAC (the reverse complement of the START codon ATG or GTG) located in the hot-spot editing window without considering any specific PAM. As a proof-of-concept, we designed gRNA04 **(Table S3)** with an ATT PAM, targeting the non-coding strand of *glnA*, where the nucleotide C in CAT, was located in position −19 for maximal editing efficiency (Figure 2A). A successful editing would lead to the elimination of the START codon, thus inactivating *glnA*.

**Figure 2.**
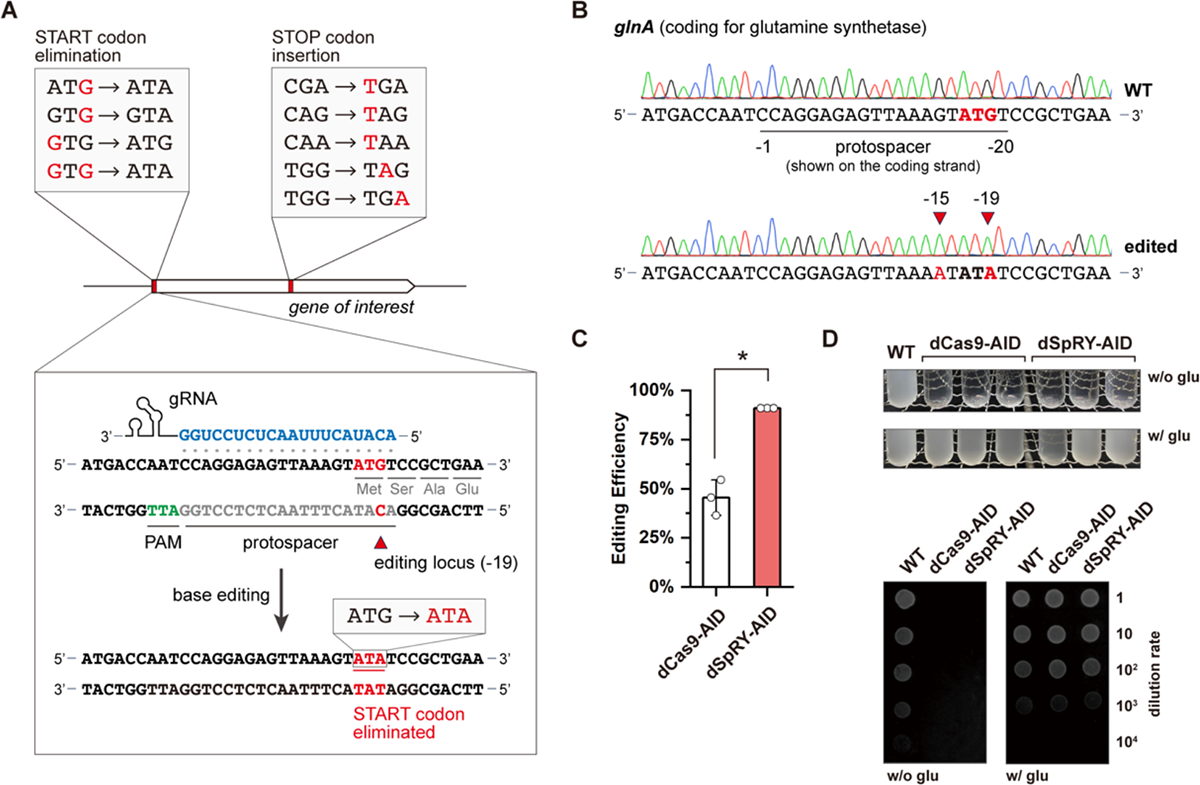
Design principle and demonstration of XSTART. **(A)** Principles of XSTART. The designed strategy, in theory, can inactivate any GOIs by designing a gRNA targeting the non-coding strand with CAT or CAC (the reverse complement of the START codon ATG or GTG) located in the hot-spot editing window without considering any specific PAM. **(B)** Sequencing results of *glnA* in the wild-type and the edited strain with XSTART. The edited loci are indicated by red arrows and highlighted in red. **(C)** Editing efficiencies of XSTART with dCas9-AID and dSpRY-AID as effectors. The results are shown as the means of three biologically independent experiments. The error bars indicate the SDs, and the differences were statistically evaluated by *t*-test (*p < 0.05). **(D)** Phenotypical evaluation of the *glnA* inactivated strain.

As a result, the START codon was successfully excluded, changing ATG to ATA (Figure 2B), and we observed an editing efficiency of 90.91% for all three individual rounds of experiments (Figure 2C). We found bystander editing at position −15 of the spacer where the G in the 5’-Untranslated region was converted to A (a C-to-T editing on the non-coding strand), but it would not influence the efficacy of our system (Figure 2A). Interestingly, we found that XSTART could also work with dCas9-AID with a non-NGG PAM for this specific target, which may result from the TGG motif next to the designed ATT PAM (Figure 2A). This agreed with a previous report using dCas9 mediated base editing to eliminate START codons in rabbit models, but a NGG or NGN PAM is compulsory (Chen et al., 2020). We compared the editing efficiencies of these two systems in *E. coli*, and found that dSpRY-AID had a significantly higher efficiency (90.91%) compared to that of dCas9-AID (45.45 ± 9.09%) (Figure 2C). This, as designed, can be explained by the released dependence on PAM and the relocation of the target nucleotide C in the preferred editing window. Next, we examined the phenotypical changes of the edited strains both in liquid medium and agar plates. As expected, the edited strains cannot grow in minimal medium without the supplement of glutamine (Figure 2D), indicating that XSTART could be a promising strategy for gene inactivation.

### Reprogramming amino acid metabolism in *E. coli* with multiplex XSTART

Next, we attempted to reprogram the amino acid metabolisms of *E. coli* with XSTART. While we already demonstrated that XSTART can inactivate *glnA* and generate a glutamine auxotrophic strain, we selected the arginine metabolism as a second target. The inactivation of *argH,* coding for the argininosuccinate lyase, would result in an arginine auxotrophic strain (Figure 3A). With gRNA05 targeting the START codon of *argH*, we obtained efficient conversion of ATG to ATA, excluding the START codon (Figure 3B), and the resulting strain can only grow in minimal medium with the supplement of arginine (Figure 3C). Then, we tried to generate a tyrosine auxotrophic strain by inactivating *tyrA*, coding for the fused chorismate mutase T or prephenate dehydrogenase, and *tyrB*, coding for the tyrosine aminotransferase, respectively (Figure 3D). Though we found successful elimination of the START codon of *tyrA*, the edited strain could still grow without tyrosine **(Figure S3)**, and similar success in genome editing but failure in phenotypical verification were observed for *tyrB* **(Figure S4)**.

**Figure 3.**
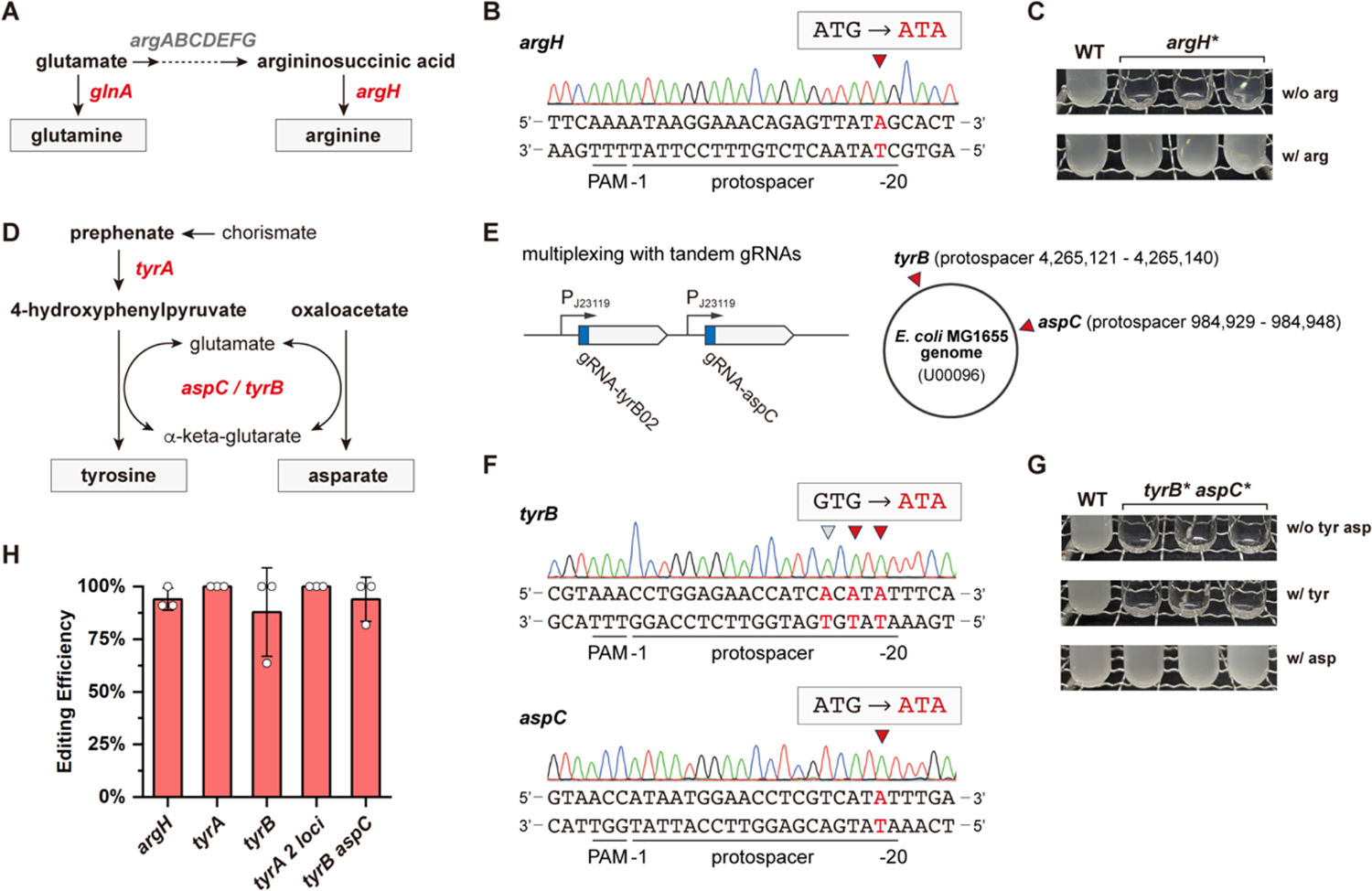
Perturbation of the amino acid metabolisms in *E. coli* with multiplex XSTART. **(A)** Schematic illustration of the glutamine and arginine metabolisms in *E. coli*. The *argH* gene encoding for the argininosuccinate lyase and *glnA* encoding for the glutamine synthetase are highlighted. **(B)** Sequencing results of *argH* edited with XSTART. **(C)** Phenotypical evaluation of *argH* inactivated strain. **(D)** Schematic illustration of the tyrosine and asparate metabolisms in *E. coli*. The *tyrA* gene encoding for the fused chorismate mutase T or prephenate dehydrogenase, the *aspC* gene encoding for the aspartate aminotransferase, and the *tyrB* gene encoding for tyrosine aminotransferase are highlighted. **(E)** Design of the tandem gRNAs (gRNA07 and gRNA09) targeting *tyrB* and *asp*C simultaneously. The tandem gRNA cassette is under the control of the constitutive promoter P_J23119_. The loci of protospacers in *tyrB* and *aspC* in the *E. coli* MG1655 genome are indicated by red arrows. **(F)** Sequencing results of *tyrB* and *aspC* edited by the multiplex XSTART. **(G)** Phenotypical evaluation of the *tyrB* and *aspC* double inactivated strain. **(H)** Editing efficiencies of XSTART in *E. coli*. The means of the editing efficiencies from three independent replicates are shown. The edited loci are indicated by red arrows and highlighted in red. Three independent isolates of the edited strain were randomly chosen for phenotypical evaluation.

We closely checked the CDS of *tyrA* and found another ATG located downstream of the original START codon **(Figure S5)**. Therefore, we hypothesized the resulting strain might contain a N-terminus truncated *tyrA* rather than an inactivated one. To check this assumption, we generated a multiplex XSTART system via assembly of tandem gRNAs (gRNA06 and gRNA08) targeting the two ATG at once. As expected, the two loci were both edited and the ATGs were removed **(Figure S5)**. But still, the resulting strain can grow without tyrosine, indicating alternative pathways supplying tyrosine for cell growth. As previously reported, *tyrB* and *aspC* (coding for aspartate aminotransferase) share similar catalytical activities, the function of which can be complemented by each other (Guzman et al., 2015). So, we deployed the multiplex XSTART for the inactivation of both genes with gRNA07 and gRNA09 (Figure 3E). We successfully obtained the strain with both genes edited (Figure 3F). Surprisingly, we found the resulting strain with the deficient *tyrB* and *aspC* cannot grow in minimal medium with or without tyrosine, but it could survive the minimal medium with the supplement of aspartate, giving an aspartate auxotrophic strain (Figure 3G). This is in line with a previous study discussing amino acid metabolisms in *E. coli* (Iwasaki et al., 2021), but the mechanism remains unclear. It was worth noticing that the editing efficiencies of XSTART were overall above 87.88% for all the above-mentioned experiments (Figure 3H), indicating the grand potential of this universal strategy. The unexpected phenotypes of the edited strains may due to the complex metabolisms of amino acids (Iwasaki et al., 2021; Miyakoshi, 2024).

### Universality of XSTART

Moreover, we selected the probiotic EcN and cyanobacterium *S. elongatus* PCC7942 to demonstrate the universality of XSTART. As EcN is a clinical isolate of *E. coli* (Lynch et al., 2022), we deployed the same system as for the model strain and targeted the *argH* gene to generate an auxotrophic EcN. The resulting strain would be useful as a probiotic chassis to harbor functional plasmids with auxotrophic markers for selection, avoiding the utilization of antibiotic resistance genes (Amrofell et al., 2023; Koh et al., 2022). By using XSTART, we managed to inactivated *argH* in EcN (Figure 4A), and the edited strain can only grow in a minimal medium with the supplement of arginine (Figure 4B).

**Figure 4.**
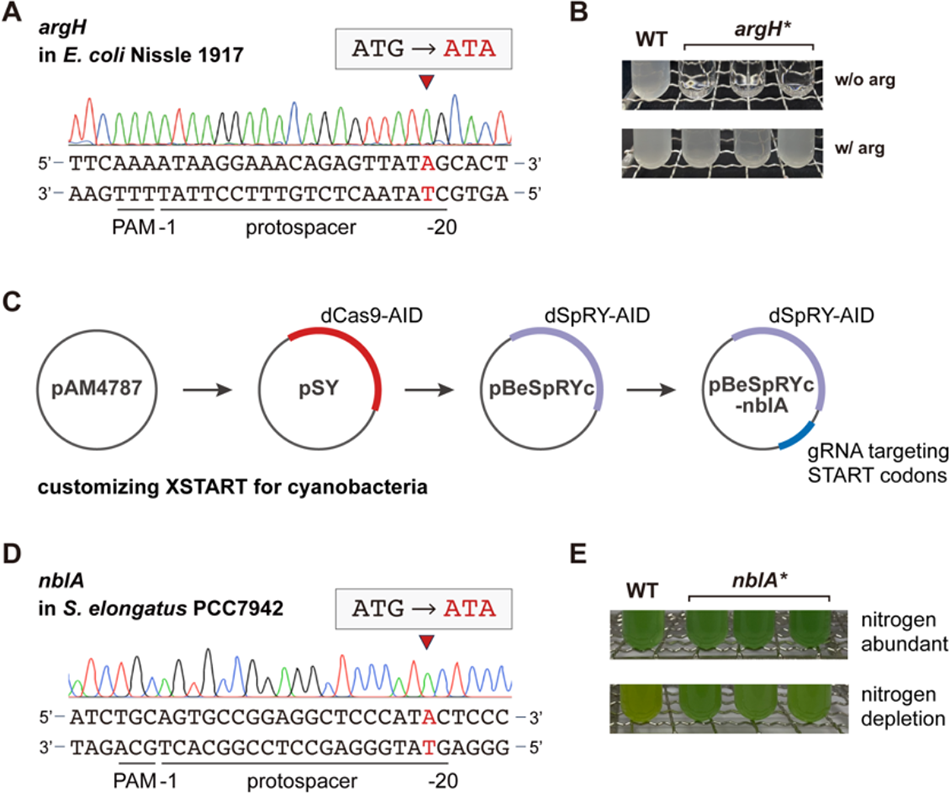
Base editing in EcN and cyanobacterium *S. elongatus* PCC7942 with XSTART. **(A)** Sequencing results of *argH* in EcN edited by XSTART with pBeSpRY-argH. **(B)** Phenotypical evaluation of the *argH* inactivated EcN. **(C)** Customization of XSTART for cyanobacteria via replacing the dCas9-AID module on pSY plasmid with the dSpRY-AID module. **(D)** Sequencing results of *nblA* in *S. elongatus* PCC7942. **(E)** Phenotypical evaluation of the *nblA* inactivated *S. elongatus* PCC7942 in nitrogen rich and depletion conditions. The edited loci are indicated by red arrows and highlighted in red. Three independent isolates of the edited strain were randomly chosen for phenotypical evaluation.

To allow XSTART work for *S. elongatus*, we generated a working plasmid via updating our previously established pSY serial plasmids with dSpRY-AID substituting dCas9-AID on the pAM4787 backbone (Figure 4C) (Chen et al., 2016; Wang et al., 2023). Then, we designed XSTART by targeting the START codon of *nblA*, coding for the phycobilisome hydrolyzing enzyme, with gRNA10 (Figure 4D). The inactivation of *nblA* in *S. elongatus* will no longer exhibit the bleaching phenotype under nitrogen-limited condition (Sendersky et al., 2014). With the customized XSTART for *S. elongatus*, we successfully inactivated *nblA* (Figure 4D), and a clear phenotype was observed with the gene-edited strain (Figure 4E). These results demonstrated the generalizability of XSTART in different strains via modular assembly of the essential modules and programming gRNAs with a straightforward design principle.

### Trade-offs between universality and precision

Unlike conventional CRISPR-Cas-based genome editing, base editing leads to unwanted off-target events. Therefore, we performed whole-genome sequencing (WGS) to analyze the off-target events caused by XSTART. Four edited strains have been selected, each with three biological independent colonies, including *glnA* edited by dCas9-AID, by dSpRY-AID, *argH* edited by dSpRY-AID, and *tryB* and *aspC* edited by dSpRY-AID (Figure 5). According to the WGS data, we found unexpected high frequency of off-target events comparing to previous reports using AID from *P. marinus* (Banno et al., 2018; Wang et al., 2023; Wei et al., 2023). All off-target editing were C-to-T or G-to-A transitions probably due to deamination, agreeing with previous reports (Rodrigues et al., 2021; Tong et al., 2019). Especially, we observed only less than 10 off-target events when inserting premature STOP codons in *S. elongatus* (Wang et al., 2023) and *Roseovarius nubinhibens* (Wei et al., 2023). The off-target events of XSTART ranged from 41 to 303 with no statistically differences among the four different editing **(Dataset)**, while the abnormally high off-target events (303) only occurred in one colony with edited *glnA* (Figure 5). To be noticed, the numbers of the off-target events between dSpRY-AID and dCas9-AID were also not statistically different (P = 0.12), while that of dSpRY-AID was slightly higher, indicating that the high frequency of off-target events resulted from the targeting of START codons rather than the released PAM dependency. Finally, we acknowledge that a proportion of these off-target events (26.2% − 53.7%, **Dataset**) led to missense mutations and early STOP codons brought by these off-target edits, while no observable influences on the designed phenotypes were identified. Taking together, we conclude a higher off-target editing caused by XSTART which might be an inevitable trade-off with the universality of this strategy.

**Figure 5.**
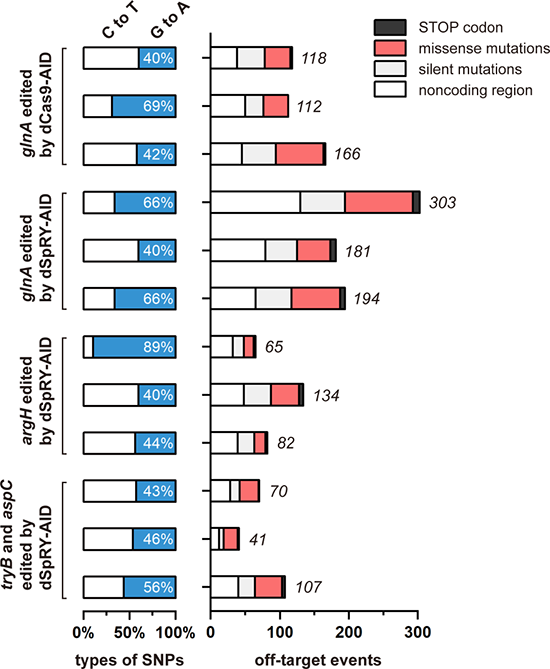
Analysis of the off-target events with whole-genome sequencing. The distribution of different types of SNPs and the consequential mutations are shown, including missense (red), nonsense (premature STOP codon, black), silent mutations (light grey), and the mutations that did not lie in the CDS (white). The analysis was performed on four differently edited strains, and three independent isolates for each strain were chosen for whole-genome sequencing.

## Discussion

Here, we report a one-for-all gene inactivation strategy, XSTART, for bacteria leveraging dSpRY-driven base editing. SpRY, the engineered Cas9 derivative, exhibited unusual advantages for developing base editors due to the independence of PAMs, thus significantly expanding the utility of base editing (Christie et al., 2023; Walton et al., 2020; Zhao et al., 2023). In return, a base-editing system using dSpRY does not cut DNAs, avoiding the cleavage of DNA coding for gRNAs. This is because SpRY can hardly distinguish the target DNA sequence from the gRNA sequence on plasmid, and we did observe mutations in the gRNA region of our working plasmids **(Figure S6)**. After releasing the PAM dependence, we designed XSTART to mutate the START codon rather than to insert STOP codons. A START codon intrinsically exists in CDS coding for the first amino acid of a protein, such as ATG or GTG. As long as one of the Gs is converted, a START codon can be eliminated. To the contrary, one must find the four specific codons for the insertion of premature STOP codons, which is sometimes difficult or impossible. Therefore, the combination of PAM-independent base editing and the principle of START codon exclusion finds a unique but wide niche for gene inactivation in bacteria.

We noticed that recent studies also recognized similar potentials of unconstrained base editing in bacteria. Two reports described a strategy by using cytosine base editor to insert STOP codons and using adenine base editor to manipulate START codons for efficient gene editing in *Bacillus subtilis* and *Pseudomonas putida*, respectively (Kozaeva et al., 2024; Xia et al., 2023). An earlier study in two rabbit models utilized dCas9-based base editor to mutate START codons, but a NGG or NGN PAM is compulsory, limiting its utility (Chen et al., 2020). Differently, our XSTART, with a single base-editing system, can be a one-for-all strategy that is capable of inactivating any GOIs by simply designing a gRNA targeting the universal START codons from the non-coding strand, without switching between different types of base editors, calculating the PAM and editing window, nor digging for the candidate codons to introduce premature STOP codons.

One essential feature is that base editing bypasses the strong “dead-or-alive” selection from DSBs, making it preferable for bacteria that are sensitive to Cas nuclease and lack efficient HR. However, without a DSB, off-target events become inevitable, as an off-target binding of base editor will not kill the bacteria anymore. This is actually true and has been carefully analyzed as reported previously in, to name a few, *S. elongatus* (Wang et al., 2023), *Streptomycetes* (Whitford et al., 2023), *Roseovarius nubinhibens* (Wei et al., 2023), *Clostridium autoethanogenum* (Seys et al., 2023) and *P. putida* (Volke et al., 2022). According to the literature and our previous work, base editors with AID showed countable off-target events and most of them resulting from the deamination (Banno et al., 2018; Wang et al., 2023; Wei et al., 2023), while C-terminus APOBEC1 (apolipoprotein B mRNA-editing enzyme) based system showed higher but acceptable frequency of off-target events (Rodrigues et al., 2021; Tong et al., 2019; Whitford et al., 2023). Though XSTART showed great capability of gene inactivation, a price of higher frequency of off-target events has been paid. We observed larger number of off-target editing using XSTART comparing to early STOP codon insertion using similar base editors, while all designed phenotypes were not influenced. This might be an inevitable trade-off for a universal and highly efficient gene editing tool under current design, where innovations with fundamental advances are on demand.

## Methods

### Strains and Media

All strains used in this study are listed in **Table S1**. *E. coli* DH5α (Takara Bio. Tech.) was used for molecular cloning to construct plasmids. *E. coli* MG1655 (CGSC#6300) strain, EcN strain and cyanobacteria *S. elongatus* PCC7942 (ATCC 33912) strain were used to test the feasibility of base editing system. EcN was a generous gift from Dr. Chun Loong Ho. Both *E. coli* strains were cultivated in Luria-Bertani (LB) medium (5 g/L yeast extract, 10 g/L tryptone, 10 g/L NaCl; solid medium with 1.5% agar) supplemented with ampicillin (150 μg/mL) or spectinomycin (60 μg/mL) when appropriate. All *E. coli* strains were cultivated at 37 °C, while MG1655 carrying plasmids with the temperature-sensitive origin of replication were grown at 30 °C. Cyanobacteria were cultivated in standard or nitrate-depleted BG-11 medium (solid medium with 1.5% agarose) at 30 °C with continuous illumination (30 – 40 µmol protons m^-2^×s^-1^). When necessary, spectinomycin (2 μg/mL) and streptomycin (2 μg/mL) were added to the medium. IPTG (0.1 mM) was added to the medium to induce base editing.

### Plasmid construction

To build the editing plasmid pBeSpRY for *E. coli*, we first chose the pKD46 plasmid with temperature-sensitive origin of replication, thus allowing plasmid curing. DNA fragments containing the *lacI*-P_trc_ inducible system, *PmCDA1* AID, *ugi* and the LVA tag were amplified from the plasmid pSY constructed in our previous study (Wang et al., 2023), and the gene *SpRY* was amplified from the plasmid pCMV-T7-SpRY-P2A-EGFP (Walton et al., 2020). The above modules were assembled to obtain the plasmid pAM4787-SpRY-AID, and then, the working plasmid pBeSpRYc was constructed by mutating SpRY with D10A and H840A. The gRNA cassette targeting specific genome locus was constructed by inverse PCR on the plasmid pTemplate as illustrated in our previous study (Wei et al., 2023), and was embedded in pBeSpRY via In-Fusion assembly, giving the base editing plasmid. To construct the base editor pBeSpRYc-nblA for *S. elongatus* PCC7942, we replaced the dCas9-AID module with dSpRY-AID in pSY plasmid, and integrated the gRNA cassette targeting *nblA* to generate the working plasmid.

All plasmids used in this study are listed in **Table S2**. All gRNA sequences designed are shown in **Table S3**, and all primers used in this study (ordered from Beijing Genomics Institute) are listed in **Table S4**. DNA fragments were amplified using PrimeSTAR Max DNA polymerase (Takara Bio. Tech.) and assembled into the vector using In-Fusion Snap Assembly Premix Kit (Takara Bio. Tech.). All plasmids were extracted using TIANprep Mini Plasmid Kit (TIANGEN Biotech.), verified by Sanger sequencing and quantified using NanoDrop One Microvolume UV-Vis Spectrophotometer (Thermo Fisher Scientific).

### Transformation of *E. coli* and *S. elongatus*

*E. coli* competent cells were prepared as described in our previous study (Li et al., 2023). In brief, freshly cultured *E. coli* cells were gently washed with pre-cooled CaCl_2_ (0.1 M), and the cells were re-suspended in 0.1 M of CaCl_2_ with 15% (v/v) of glycerol. The competent cells were stored at −80 °C before use. The base editing working plasmid (60 - 80 ng) was added into 100 µL MG1655 or Nissle 1917 competent cells, while the same volume of ddH_2_O was added for control group. The mixture was placed on ice for 20 min, heated shock at 42 °C for 1 min, and transferred to 3 mL fresh LB media with 3 µL ampicillin. After cultivation at 30 °C for 1 h, cells were induced for 3 h with 0.1 mM IPTG, and plated on LB agar plates with ampicillin to obtain transformants. To transform the cyanobacterium, *S. elongatus* PCC7942 cells were cultivated to OD_730_ of 0.5 - 0.7 and then 15 mL of culture was centrifuged at 5,000 rpm for 15 min at 24 °C.

After washing with fresh BG-11, the cells were re-suspended in 300 µL BG-11, and 2 µg plasmid DNA was added and incubated overnight at 30 °C under dark light. The samples were then transferred to a culture tube with appropriate antibiotics. IPTG was added to induce base editing after 24 h of cultivation at 30 °C, 120 rpm. After induction culture for 48 h, cell suspensions were plated on BG-11 plates with appropriate antibiotics, and single colonies were randomly selected for analysis.

### Colony PCR and Sanger Sequencing

Transformants were randomly selected for colony PCR. The single colony was suspended in 20 µL ddH_2_O, cleaved at 100 °C for 10 min, and 1 µL of supernatant was taken as template DNA. The 20-µL PCR reaction system consisted of 10 µL PrimeSTAR Max DNA polymerase, 0.5 µL positive and negative primers, 1 µL template DNA, and 8 µL ddH_2_O. The PCR products were checked by Sanger sequencing.

### Plasmid Curing

After base editing, the edited pure strains were inoculated in LB medium and cultivated overnight at 40 °C. The cells were streaked on LB agar plate for analysis. 6 - 8 single colonies were randomly selected to check the presence of plasmids, while the wild type stain (without plasmids) and the working plasmid (with plasmid) were employed as controls. To further verify that the plasmid had been eliminated, individual colony was inoculated to LB solid media with or without ampicillin. The plasmid was cured when the strain can grow without antibiotics but cannot grow with antibiotics

### Phenotypic Verification

Both wild type and the engineered *E. coli* strains were cultivated in M9 minimal medium (15.14 g/L Na_2_HPO_4_·12H_2_O, 3.0 g/L KH_2_PO_4_, 0.5 g/L NaCl, 1.0 g/L NH_4_Cl, 0.241 g/L MgSO_4_, 0.011 g/L CaCl_2_ and 4 g/L glucose solution; solid medium with 1.5% Agar) or in M9 medium supplied with specific L-amino acids to check the growths of the engineered strains. The working concentrations of amino acids added to M9 minimal medium are as follows: 5 mM L-glutamine, 1 mM L-tyrosine, 0.4 g/L L-arginine and 0.4 g/L L-aspartate.

### Whole-Genome Sequencing

Whole-genome sequencing was carried out to assess off-target events in modified *E. coli* strains, following previously established method in our lab (Wei et al., 2023). *E. coli* cells at the exponential phase were harvested in volumes of 30 – 50 mL. Following the extraction of the genomic DNA of the sample, qualified DNA samples were randomly broken into 350 – 500 bp fragments by Covaris. After library construction, sequencing was performed by Illumina HiSeq instrument. After the process of quality filtering of the original sequencing data, Clean Reads were compared to the reference genome (U00096 in this study) using BWA, and the results were analyzed with Qualimap software. The results can be found in the NCBI SRA with the accession number PRJNA1122099.

## Supporting Information

Summary of strains, plasmids, primers and gRNA sequences; summary of the editing efficiency; and figures supporting the main results.

## Acknowledgments

This work was supported by the National Natural Science Foundation of China (22278246, U20A20146 and 22378233), the Department of Science and Technology of Shandong Province (2022HWYQ-017), the Natural Science Foundation of Shandong Province (ZR2021ME066) and the Qilu Young Scholar Program of Shandong University (to P.-F.X.), and the Taishan Scholars Project of Shandong Province (NO. tstp20230604).

