## Supplementary material for "One-for-all gene inactivation via PAM-independent base editing in bacteria"

**Table S1.** Strains used in this study

| strains | description | sources |
|---|---|---|
| *E. coli* DH5α | Commercial *E. coli* strain for molecular cloning | Takara Bio. Tech. |
| *E. coli* MG1655 | Wild type strain for the evaluation of the system | Lab stock (CGSC#6300) |
| *E. coli* Nissle 1917 | Clinical *E. coli* isolate | Ho lab (Ho et al., 2018) |
| *S. elongatus* PCC7942 | Model cyanobacterium strain | Lab stock (ATCC33912) |
| *E. coli* LX01 | MG1655, *glnA* inactivation by dSpRY-AID system with ATG START codon mutating to ATA | This study |
| *E. coli* LX02 | MG1655, *glnA* inactivation by dCas9-AID system with START codon ATG mutating to ATA | This study |
| *E. coli* LX03 | MG1655, *argH* inactivation by dSpRY-AID system with START codon ATG mutating to ATA | This study |
| *E. coli* LX04 | MG1655, *aspC* and *tyrB* inactivated by dSpRY-AID system with START codons ATG and GTG mutating to ATAs | This study |
| *E. coli* LX05 | MG1655, *tyrA* inactivated by dSpRY-AID system with ATGG mutating to ATAA | This study |
| *E. coli* LX06 | MG1655, *tyrA* inactivated by dSpRY-AID system with two ATG codons mutating to ATAs | This study |
| *E. coli* LX07 | MG1655, *tyrB* inactivated by dSpRY-AID system with GTG mutating to ATA | This study |
| *E. coli* LX08 | Nissle 1917, *argH* inactivation by dSpRY-AID system with START codon ATG mutating to ATA | This study |
| *S. elongatus* LX01 | PCC7942, *nblA* inactivation by dSpRY-AID system with START codon ATG mutating to ATA | This study |

**Table S2.** Plasmids used in this study

| Name | Description | References |
|---|---|---|
| pAM4787 | *ColE1 ori*, partial sequence from pANS | Lab stock |
| pSY | *ColE1 ori*, *SmR*, dSpRY-PmCDA1-*ugi* | Lab stock |
| pCMV-T7-SpRY-P2A-EGFP | *ColE1 ori*, *bla*, *SpRY* | Addgene #275392 (Walton et al., 2020) |
| pKD46 | *oriR101*, *bla* | Lab stock (Datsenko and Wanner, 2000) |
| pTemplate | *pUC ori*, gRNA, *bla* | Lab stock |
| pAM4787-SpRY-AID | *ColE1 ori*, *SmR*, SpRY-PmCDA1-*ugi* | This study |
| pBeSpRYc | pAM4787*, SmR*, dSpRY-PmCDA1-*ugi* | This study |
| pBeSpRY | *oriR101*, *bla*, *trc* promoter, dSpRY-PmCDA1-*ugi* | This study |
| pBeCas9 | *oriR101*, *bla*, *trc* promoter, dCas9-PmCDA1-*ugi* | This study |
| pgRNA01 | pTemplate, gRNA-*glnA*-stop-15 | This study |
| pgRNA02 | pTemplate, gRNA-*glnA*-stop-19 | This study |
| pgRNA03 | pTemplate, gRNA-*glnA*-stop-20 | This study |
| pgRNA04 | pTemplate, gRNA-*glnA* | This study |
| pgRNA05 | pTemplate, gRNA-*argH* | This study |
| pgRNA06 | pTemplate, gRNA-*tyrA* | This study |
| pgRNA07 | pTemplate, gRNA-*tyrB*02 | This study |
| pgRNA08 | pTemplate, gRNA-*tyrA* 2 loci | This study |
| pgRNA09 | pTemplate, gRNA-*aspC* | This study |
| pgRNA10 | pTemplate, gRNA-*nblA* | This study |
| pBeSpRY-glnA15 | pBeSpRY, gRNA01 | This study |
| pBeSpRY-glnA19 | pBeSpRY, gRNA02 | This study |
| pBeSpRY-glnA20 | pBeSpRY, gRNA03 | This study |
| pBeCas9-glnA15 | pBeCas9, gRNA01 | This study |
| pBeCas9-glnA19 | pBeCas9, gRNA02 | This study |

| | | |
|---|---|---|
| pBeCas9-glnA20 | pBeCas9, gRNA03 | This study |
| pBeSpRY-glnA | pBeSpRY, gRNA04 | This study |
| pBeCas9-glnA | pBeCas9, gRNA04 | This study |
| pBeSpRY-tyrA | pBeSpRY, gRNA06 | This study |
| pBeSpRY-tyrA-2loci | pBeSpRY, gRNA06, gRNA08 | This study |
| pBeSpRY-argH | pBeSpRY, gRNA05 | This study |
| pBeSpRY-aspC | pBeSpRY, gRNA09 | This study |
| pBeSpRY-aspC-tyrB | pBeSpRY, gRNA07, gRNA09 | This study |
| pBeSpRYc-nblA | pBeSpRYc, gRNA10 | This study |

**Table S3.** gRNA sequences used in this study

| gRNA | Target | Strand [a] | PAM | Protospacer |
|---|---|---|---|---|
| gRNA01 | *glnA* | C | NCT | AAGAACAGCACGTCACTATC |
| gRNA02 | *glnA* | C | NTC | ACAGCACGTCACTATCCCTG |
| gRNA03 | *glnA* | C | NCA | CAGCACGTCACTATCCCTGC |
| gRNA04 | *glnA* | N | NTT | ACATACTTTAACTCTCCTGG |
| gRNA05 | *argH* | N | NTT | CCATAACTCTGTTTCCTTAT |
| gRNA06 | *tyrA* | N | NAC | CCATAATAAACCTCTTAAGC |
| gRNA07 | *tyrB* | N | NTT | ACACGCGATGGTTCTCCAGG |
| gRNA08 | *tyrA* | N | NGA | ACATAGATGCCTCGCGCTCC |
| gRNA09 | *aspC* | N | NGT | ACATGACGAGGTTCCATTAT |
| gRNA10 | *nblA* | N | NCA | GCATGGGAGCCTCCGGCACT |

[a] C stands for coding strand, and N stands for non-coding strand.

**Table S4.** Primers used in this study

| Primer | Sequence |
|---|---|
| **Primers for cloning** | |
| XIA-LX-247 | CAGCTGGGAGGCGACGGAGGTGGAGGAGGTTCTGGAGG |
| XIA-LX-223 | ATGCTGTACTTCTTGTCCATCATGGTCTGTTTCCTGTGTG |
| XIA-LX-246 | CCAGAACCTCCTCCACCTCCGTCGCCTCCCAGCTGAGACA |
| XIA-LX-225 | CACACAGGAAACAGACCATGATGGACAAGAAGTACAGCATCGGCCT |
| XIA-LX-263 | GAAGTACAGCATCGGCCTGGCTATCGGCACCAACTCTGTG |
| XIA-LX-264 | AAGCTCTGAGGCACGATGGCGTCCACATCGTAGTCGGACA |
| XIA-LX-265 | CCGACTACGATGTGGACGCCATCGTGCCTCAGAGCTTTCT |
| XIA-LX-266 | ACAGAGTTGGTGCCGATAGCCAGGCCGATGCTGTACTTCT |
| XIA-LX-295 | ATTCGAAACCGGTATCCGCAGGTGGCACTTTTCGGGGAAA |
| XIA-LX-298 | TGTCTTGCGTCTCTGTCGACGGGTATGGACAGTTTTCCCT |
| XIA-LX-293 | TCCCCGAAAAGTGCCACCTGCGGATACCGGTTTCGAATTG |
| XIA-LX-297 | AGGGAAAACTGTCCATACCCGTCGACAGAGACGCAAGACA |
| XIA-LX-350 | CCATAATAAACCTCTTAAGCGTTTTAGAGCTAGAAATAGC |
| XIA-LX-351 | GCTTAAGAGGTTTATTATGGGCTAGCATTATACCTAGGAC |
| XIA-LX-352 | ACATACTTTAACTCTCCTGGGTTTTAGAGCTAGAAATAGC |
| XIA-LX-353 | CCAGGAGAGTTAAAGTATGTGCTAGCATTATACCTAGGAC |
| XIA-LX-360 | AGCGGATTTGAACGTTGCGAATCCTTGACAGCTAGCTCAG |
| XIA-LX-361 | CCAGCTCGGTCTAGATTGCTCCTTTGAGTGAGCTGATACC |
| XIA-LX-362 | CTGAGCTAGCTGTCAAGGATTCGCAACGTTCAAATCCGCT |
| XIA-LX-363 | CGGTATCAGCTCACTCAAAGGAGCAATCTAGACCGAGCTGG |
| XIA-LX-396 | AAGAACAGCACGTCACTATCGTTTTAGAGCTAGAAATAGC |
| XIA-LX-397 | GATAGTGACGTGCTGTTCTTGCTAGCATTATACCTAGGAC |
| XIA-LX-398 | ACAGCACGTCACTATCCCTGGTTTTAGAGCTAGAAATAGC |
| XIA-LX-399 | CAGGGATAGTGACGTGCTGTGCTAGCATTATACCTAGGAC |
| XIA-LX-400 | CAGCACGTCACTATCCCTGCGTTTTAGAGCTAGAAATAGC |
| XIA-LX-401 | GCAGGGATAGTGACGTGCTGGCTAGCATTATACCTAGGAC |
| XIA-LX-387 | CATCCGGCTCGTATAATGTG |
| XIA-LX-411 | AATGGACAACTCGCTCCGTC |
| XIA-LX-364 | GACGGAGCGAGTTGTCCATT |

| | |
|---|---|
| XIA-LX-389 | CACATTATACGAGCCGGATG |
| XIA-LX-446 | CCATAACTCTGTTTCCTTATGTTTTAGAGCTAGAAATAGC |
| XIA-LX-447 | ATAAGGAAACAGAGTTATGGGCTAGCATTATACCTAGGAC |
| XIA-LX-422 | ACATGACGAGGTTCCATTATGTTTTAGAGCTAGAAATAGC |
| XIA-LX-423 | ATAATGGAACCTCGTCATGTGCTAGCATTATACCTAGGAC |
| XIA-LX-453 | ACACGCGATGGTTCTCCAGGGTTTTAGAGCTAGAAATAGC |
| XIA-LX-454 | CCTGGAGAACCATCGCGTGTGCTAGCATTATACCTAGGAC |
| XIA-LX-404 | ACATAGATGCCTCGCGCTCCGTTTTAGAGCTAGAAATAGC |
| XIA-LX-405 | GGAGCGCGAGGCATCTATGTGCTAGCATTATACCTAGGAC |
| XIA-LX-406 | GGTATCAGCTCACTCAAAGGTTGACAGCTAGCTCAGTCCT |
| XIA-LX-407 | AACTGTCCATACCCGTCGACGAGAGCGTTCACCGACAAAC |
| XIA-LX-408 | AGGACTGAGCTAGCTGTCAACCTTTGAGTGAGCTGATACC |
| XIA-LX-409 | GTTTGTCGGTGAACGCTCTCGTCGACGGGTATGGACAGTT |
| XIA-LX-491 | GCATGGGAGCCTCCGGCACTGTTTTAGAGCTAGAAATAGC |
| XIA-LX-492 | AGTGCCGGAGGCTCCCATGCGCTAGCATTATACCTAGGAC |
| **Primers for verification** | |
| XIA-LX-258 | CTATGAGAAGCTGAAGGGCT |
| XIA-LX-273 | CATCCGGCTCGTATAATGTG |
| XIA-LX-274 | CGGTTCTTGTCGCTTCTGGT |
| XIA-LX-299 | TCGTTCTCATGGCTCACGCA |
| XIA-LX-300 | ATAAGGGCGACACGGAAATG |
| XIA-LX-364 | GACGGAGCGAGTTGTCCATT |
| XIA-LX-375 | CCTACCTACGTAACGGACTA |
| XIA-LX-391 | GACAGCTTATCATCGACTGC |
| XIA-LX-412 | CTCTCATCATACGCAGTGTG |
| XIA-LX-410 | ATGGTTCGTTCTCATGGCTC |
| XIA-LX-275 | AAGCAATCTAGACCGAGCTG |
| XIA-LX-354 | ACTGATGCCAGATCGACCAG |
| XIA-LX-355 | CAGCAATTAACGCTATGCGC |
| XIA-LX-356 | AGGCCAACATAGATGCCTCG |
| XIA-LX-357 | ACCTTCGTATTGGGTGGAGG |
| XIA-LX-358 | GAACGTACCGGATTGTTGGA |

| | |
|---|---|
| XIA-LX-359 | CAATCGAGGAGCCGTCAAAC |
| XIA-LX-384 | CCACCTTCATATTGGGTGGA |
| XIA-LX-385 | GAACGTACCGGATTGTTGGA |
| XIA-LX-386 | CAATCGAGGAGCCGTCAAAC |
| XIA-LX-418 | CTTATCCCGGATTCTCAGGA |
| XIA-LX-448 | GGATATCTTCGGCGTCGCTT |
| XIA-LX-449 | CGCGAAAGCAGAACAACTGA |
| XIA-LX-450 | CTCTTCTGCGGTTAACACGC |
| XIA-LX-419 | CAGTTAAGCCCTTCCATCGG |
| XIA-LX-420 | AGGGTCGCGATGAAATACGT |
| XIA-LX-421 | CGGCTTGCAGTTGTGGAATA |
| XIA-LX-426 | GTAGGTTCAGACGACACCGT |
| XIA-LX-427 | TGCGAGCACGTTTGTCATTG |
| XIA-LX-428 | CAGCCTTTTTCACGCTGGTC |
| XIA-LX-413 | CGATGGTGCGCATGATAACG |
| XIA-LX-493 | ACTGCGCATTTTCGTGACAC |
| XIA-LX-259 | AATGCTGCTGCCTCTACTTC |
| XIA-LX-260 | GTGTACCAGTTGCCAAGCCA |
| XIA-LX-204 | CGAGATAGCAGTATTGACGG |

**Table S5.** Base editing efficiency in this study

| Target | gRNA | Rounds of independent replicates | Number of selected colonies | Editing efficiency % | Colonies with pure edits | Pure colonies after one-round of segregation |
|---|---|---|---|---|---|---|
| **Edited by dSpRY-AID** | | | | | | |
| *glnA* | gRNA01 | 4 | 44 | 59.09 ± 22.88 | 0 | Yes |
| *glnA* | gRNA02 | 4 | 44 | 100 ± 0 | 12 | Yes |
| *glnA* | gRNA03 | 4 | 44 | 95.45 ± 9.09 | 1 | Yes |
| *glnA* | gRNA04 | 3 | 33 | 90.91 ± 0 | 0 | Yes |
| *argH* | gRNA05 | 3 | 33 | 93.94 ± 5.25 | 1 | Yes |
| *tyrA* | gRNA06 | 3 | 33 | 100 ± 0 | 20 | Yes |
| *tyrA* 2 loci | gRNA06, gRNA08 | 3 | 33 | 100 ± 0 | 29 | Yes |
| *tyrB* | gRNA07 | 3 | 33 | 87.88 ± 20.99 | 2 | Yes |
| *aspC-tyrB* | gRNA07, gRNA09 | 3 | 33 | 93.94 ± 10.50 | 1 | Yes |
| **Edited by dCas9-AID** | | | | | | |
| *glnA* | gRNA01 | 3 | 33 | 0 | 0 | Yes |
| *glnA* | gRNA02 | 3 | 33 | 0 | 0 | Yes |
| *glnA* | gRNA03 | 3 | 33 | 0 | 0 | Yes |
| *glnA* | gRNA04 | 3 | 33 | 45.45 ± 9.09 | 0 | Yes |

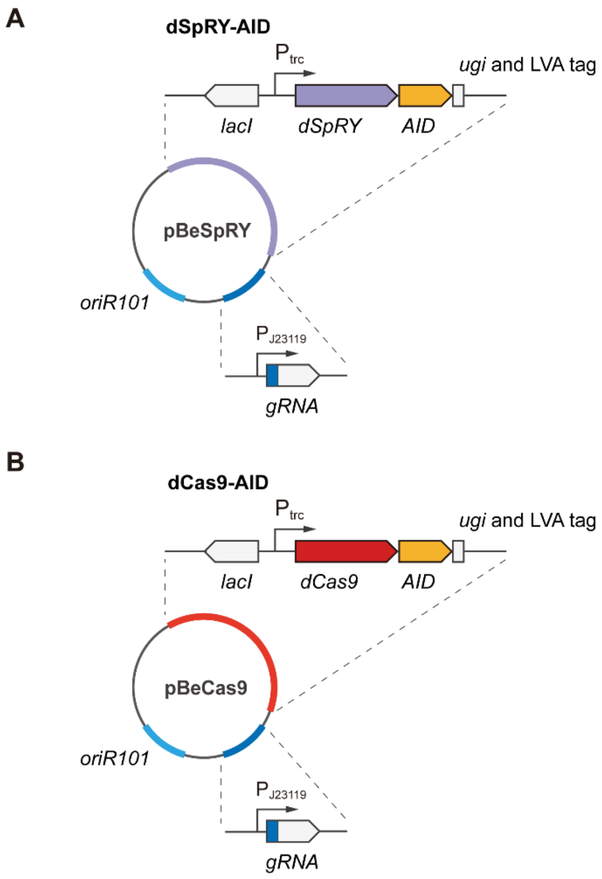


**Figure S1.** Design of the working plasmids pBeSpRY-gRNA and pBeCas9-gRNA. (A) pBeSpRY plasmid contains the dSpRY-AID module carrying *dSpRY*, *AID*, *ugi* and LVA tag driven by a *lacI-*$P_{trc}$ inducible system. The gRNA cassette is under the control of the constitutive promoter $P_{J23119}$ (B) pBeCas9 plasmid is similarly designed with dCas9-AID as the effecter, containing *dCas9*, *AID*, *ugi* and LVA tag under the control of *lacI-*$P_{trc}$ inducible system.

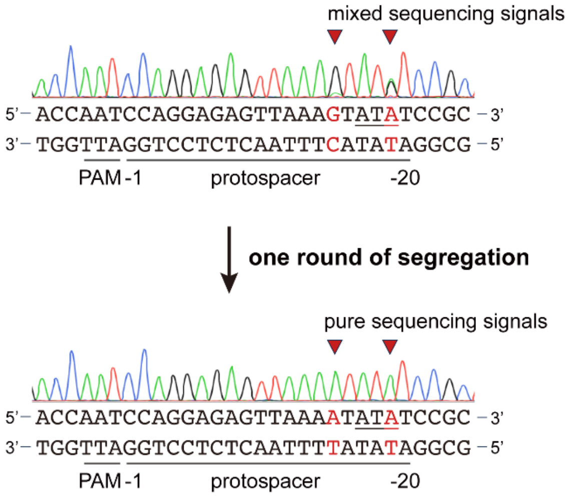


**Figure S2.** Sequencing results of the edited strain with mixed signals and the sequencing results of the pure edited strain after one more round of segregation. The edited nucleotides are indicated by red arrows and highlighted in red.

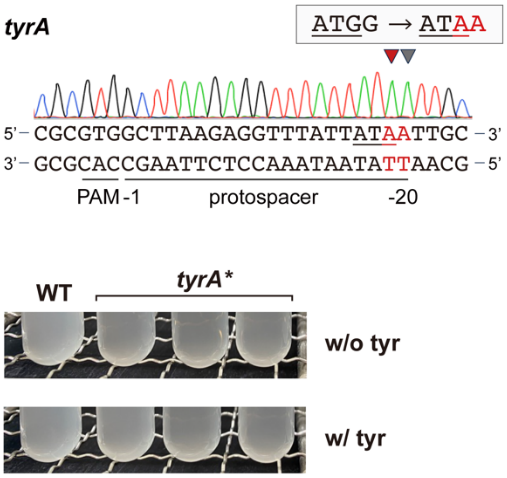


**Figure S3.** Sequencing result of *tyrA* edited by XSTART, and the phenotypical evaluation of the *tyrA* inactivated strain. The edited nucleotides are indicated by red (intended editing) and grey (bystander editing) arrows and highlighted in red. The wild type strain and three randomly picked clonies carrying designed edits in *tyrA* were cultured in minimal medium with and without 1 mM L-tyrosine.

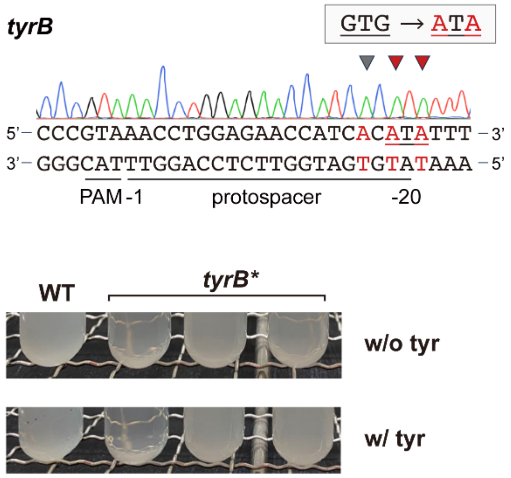


**Figure S4.** Sequencing analysis of *tyrB* edited by XSTART, and the phenotypical evaluation of the *tyrB* inactivated strain. The edited nucleotides are highlighted in red and the edited loci are indicated by red arrows. The wild type strain and three randomly picked clones carrying desired edits in *tyrB* were cultured in minimal medium with and without 1 mM L-tyrosine.

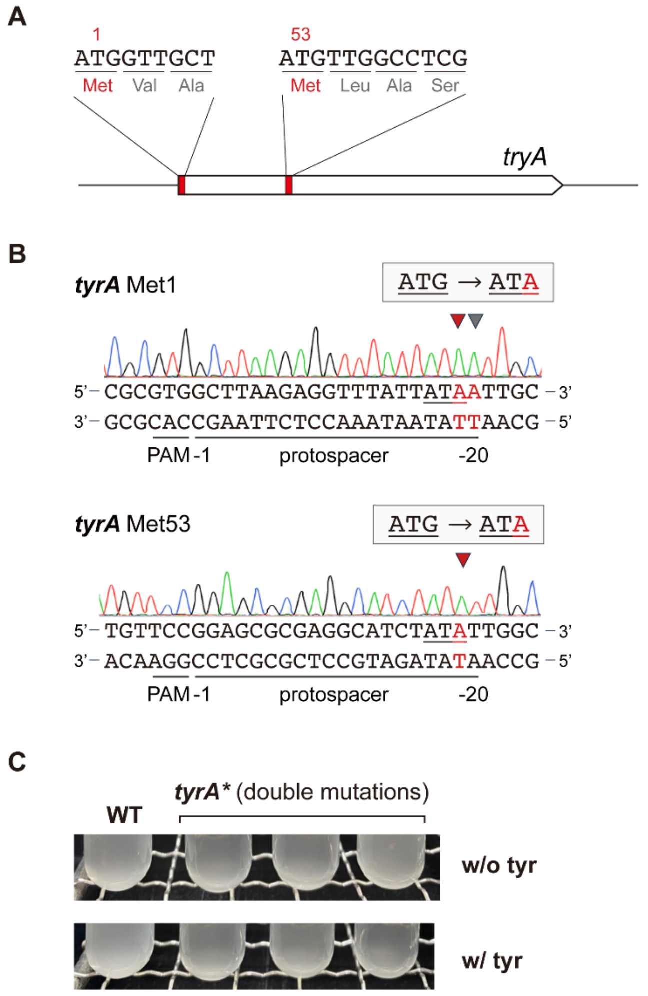


**Figure S5.** Evaluation of a multiplex XSTART system targeting *tyrA* with tandem gRNAs. (A) The two ATG targets coding for Met1 and Met53 in *tyrA*. (B) Sequencing of the two edited loci with XSTART. The edited nucleotides are highlighted in red and indicated by red arrow. Grey arrow indicates bystander editing. (C) The phenotypical evaluation of the *tyrA* inactivated strain. The wild type strain and three randomly picked clones carrying designed double-edits in *tyrA* were cultured in minimal medium with and without 1 mM L-tyrosine.

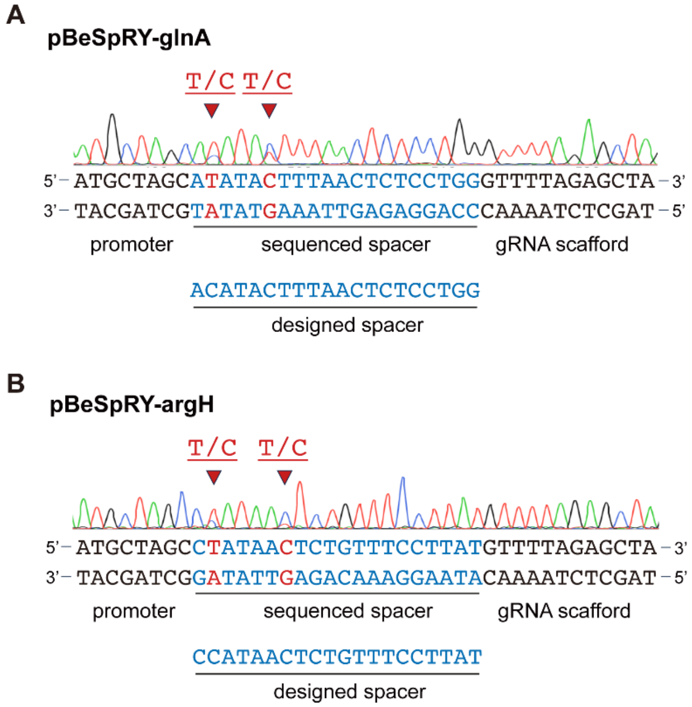


**Figure S6.** The sequencing results of the gRNA cassette on pBeSpRY-glnA (A) and pBeSpRY-argH (B). The designed spacer is highlighted in blue and the mixed sequencing signals of mutated nucleotides are highlighted with red arrows and red font.
